# Fundamental role for the creatine kinase pathway in protection from murine colitis

**DOI:** 10.1101/2023.06.07.544110

**Authors:** Caroline HT Hall, Jordi M Lanis, Alexander S Dowdell, Emily M Murphy, Sean P Colgan

## Abstract

Inflammatory diseases of the digestive tract, including inflammatory bowel disease (IBD), cause metabolic stress within mucosal tissue. Creatine is a key energetic regulator. We previously reported a loss of creatine kinases (CKs) and the creatine transporter expression in IBD patient intestinal biopsy samples and that creatine supplementation was protective in a dextran sulfate sodium (DSS) colitis mouse model. In the present studies, we evaluated the role of CK loss in active inflammation using the DSS colitis model. Mice lacking expression of CKB/CKMit (CKdKO) showed increased susceptibility to DSS colitis (weight loss, disease activity, permeability, colon length and histology). In a broad cytokine profiling, CKdKO mice expressed near absent IFN-γ levels. We identified losses in IFN-γ production from CD4^+^ and CD8^+^ T cells isolated from CKdKO mice. Addback of IFN-γ during DSS treatment resulted in partial protection for CKdKO mice. We identified basal stabilization of the transcription factor hypoxia-inducible factor (HIF) in CKdKO splenocytes and pharmacological stabilization of HIF resulted in reduced IFN-γ production by control splenocytes. Thus, the loss of IFN-γ production by CD4^+^ and CD8^+^ T cells in CKdKO mice resulted in increased colitis susceptibility and indicates that CK is protective in active mucosal inflammation.

## Introduction

The intestinal mucosa is a complex environment where in the healthy state, absorption of nutrients occurs while maintaining a barrier to avoid inflammation due to the multitude of antigens present in the lumen. Maintenance of this homeostasis is an energy intensive process which requires regulation of epithelial cell barrier function as well as trafficking of cells into and out of the mucosal tissue^1^.

The creatine-phosphocreatine cycle mediates an essential mechanism of energy distribution in all cells. Creatine serves as an energetic buffer by shuttling high energy N-phosphoryl groups as phosphocreatine that can be used to produce ATP in times and locations of increased energy usage^2, 3^. Phosphocreatine diffuses more readily than ATP^4^, allowing for the distribution of energy to areas of increased need, such as those with active actin turnover^5^. Regulation of this process is mediated by adequate creatine uptake by the creatine transporter (CRT) and by reversible phosphorylation by creatine kinases (CK)^2, 6, 7^. There are cell specific CK isoforms which are found in defined cellular locations. The mitochondrial form (CKMit) is found in the outer mitochondrial compartment and is expressed in most tissues and selective expression of the muscle type (CKM) and brain type (CKB)^6^ in different cell types. Among cells that are relevant to the intestinal mucosal environment, CKB is expressed in lymphocytes and increases in expression were found in lymphocyte activation^8^. Recently, the role of CRT and CKB have been further clarified in CD8^+^ T cells with CRT loss resulting in reduced CD8^+^ T effector viability and CKB loss resulting in loss of IFN-γ and TNF-α producing lymphocytes^9^. Additionally, CKs have been implicated in actin polymerization that is required for phagocytosis in macrophages^10^. Finally, we have previously identified that CKM, CKMit and CKB are constitutively expressed intestinal epithelial cells (IECs), where expression was enhanced in the setting of hypoxia^11^.

Ongoing inflammatory responses are associated with shifts in tissue metabolism that can fundamentally influence tissue function^12–14^. A recent analysis of the literature revisited the idea that deficiencies in energy homeostasis was associated with IBD (the original “starved gut hypothesis”^15^) and concluded that disturbances in energy balance occur at multiple levels in IBD, including dysregulation of the creatine-phosphocreatine cycle^16^. Indeed, our own work identified a key role for the creatine pathway in intestinal disease. By evaluating transcriptional expression in IBD patient biopsy samples, we found dysfunctional expression of CKM, CKB, CKMit and CRT in the setting of both Crohn’s disease and ulcerative colitis^11, 17^. Moreover, we have shown that creatine supplementation was protective in TNBS and DSS colitis mouse models with preservation of colonic histologic structure and improved symptom scoring^11^.

### Additionally, have previously identified dysfunctional epithelial cell activity such as wound healing and barrier function with the loss of CRT^17^

Given mucosal loss of creatine machinery in the intestines of inflammatory bowel disease patients, we sought to evaluate the implications of CK loss on colitis susceptibility. For these purposes, we made use of mice with global loss of CKB and CKMit in a DSS colitis model. We identified significantly worsened colitis in CKdKO mice compared to control mice by disease activity, intestinal permeability and histologic scoring. Surprisingly, analysis of CKdKO mice showed decreased pro-inflammatory cytokines in the intestinal tissue, most notably nearly absent expression of IFN-γ. Profiling transcription factor expression revealed the prominent induction of hypoxia inducible factor (HIF) in CKdKO splenocytes. Further analysis revealed that pharmacological stabilization of HIF reduces IFN-γ production by lymphocytes independent of CK expression. Together these findings implicate CK loss in increased colitis susceptibility through a mechanism involving HIF-dependent repression of IFN-γ production.

## Results

### CKdKO mice have increased disease severity in DSS colitis

CKB and CKMit are important regulators of immune cell activity and are downregulated in IBD patient samples^9, 11^. Using CKB and CKMit global deletion mice (CKdKO), we conducted DSS colitis experiments using 1.5% DSS in drinking water for 5 days followed by a 2-day washout observation period. CKdKO mice showed increased weight loss and disease activity scoring compared to control mice (which includes measurement of weight loss, bleeding and stool consistency)^18^ (Fig 1A and B). The CKdKO mice also manifested increased intestinal permeability as measured by FITC-dextran permeability assay (Fig 1C, p< 0.05), which is commonly associated with pronounced colitic disease.

**Figure 1.**
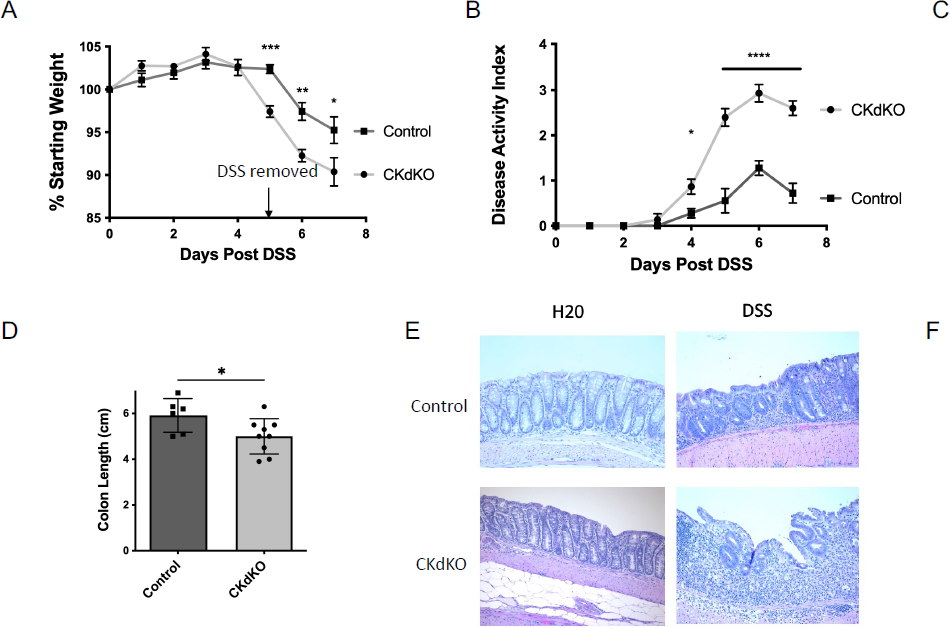
CKdKO and control mice were exposed to 1.5% DSS in their drinking water for 5 days followed by a 2-day recovery period. **a** Weights and **b** Disease Activity Index (weight change, stool consistency and bleeding score) were obtained daily. **c** On day 7, mice were gavaged with FITC-dextran 4 kD and plasma was obtained at 4 hours. **d** Colon length was measured from cecum to distal colon. **e, f** H & E of colonic tissue were compared and scored (see methods section). n=6-12 mice per group. Statistical differences were analyzed by 2way ANOVA (**a, b)**, unpaired t test (**c, d**) and one-way ANOVA (**f**). * P&lt;0.05, ** P&lt;0.01, *** P&lt;0.001, **** P&lt;0.0001

In addition to the above functional findings, the CKdKO mice had decreased colon length with DSS treatment relative to controls, which commonly is associated with increased colonic inflammation and swelling (Fig 1D, p< 0.05). Histopatholologic examination of colonic tissue revealed a profound increase in inflammation within the CKdKO cohort of mice (Fig 1E), including consistently increased inflammatory cell infiltrate, epithelial damage and nearly complete crypt loss. These histologic changes were reflected in a profound increase in histologic scores within the CKdKO cohort of mice compared to controls (Fig 1F, p< 0.001).

Taken together these results reveal a significant increase in overall colitis disease severity in the absence of CKB and CKMit.

### Prominent loss of IFN-γ expression in colitic CKdKO mice

To gain insight into mechanisms of damage, colonic tissue samples from colitic wild-type and CKdKO mice were evaluated by cytokine protein profiling. As expected, the pro-inflammatory cytokine, IFN-γ, was elevated in control DSS colitis tissues. However unexpectedly, we identified a near complete absence of IFN-γ in the tissue of DSS CKdKO mice (Fig 2A, p<0.01). While IFN-γ is thought to function of as pro-inflammatory mediator, it also has important homeostatic functions. IFN-γ can attenuate tissue injury by a number of mechanisms including inhibiting proteases and coagulation factors^19^ and reduced neutrophil and monocyte infiltration^20^. Interestingly we also identified an absence of other inflammatory cytokines, such as IL-6, Il-22 and IL-17F, in DSS CKdKO tissue compared to DSS control tissue (Supp Fig 1A), although none of these reductions were as significant as that seen in IFN-γ.

**Figure 2.**
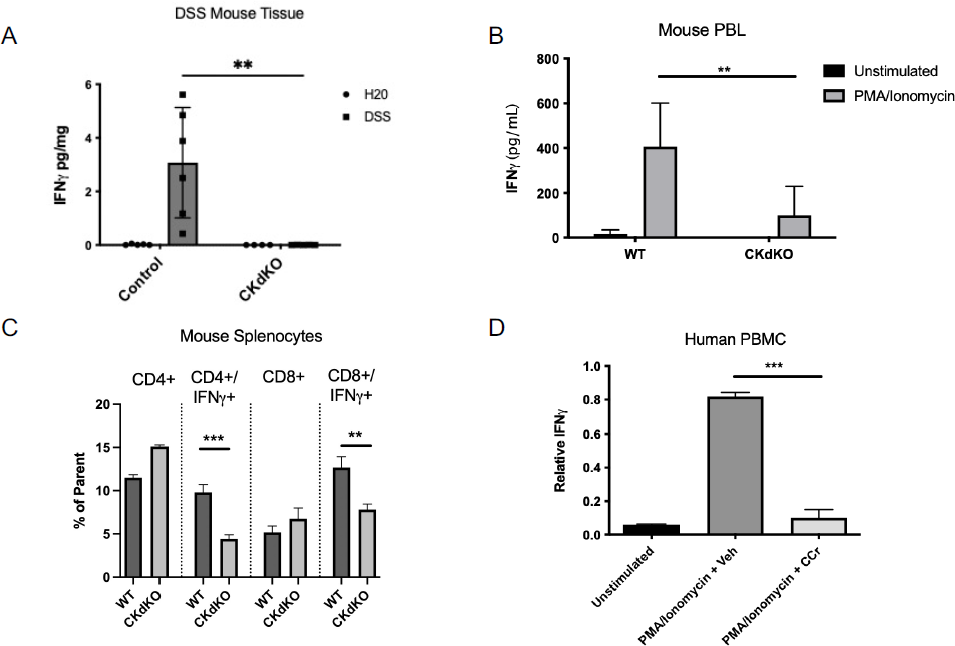
**a** Total colonic tissue samples from DSS experiment were homogenized and protein homogenate was evaluated using IFN-γ mesoscale analysis (n=5-6). **b** PBL were isolated from wildtype and CKdKO mice and stimulated with PMA and ionomycin for 18 hours followed by supernatant collection and analysis by IFN-γ ELISA (n=4-6). **c** Splenocytes from wildtype and CKdKO mice were stimulated with PMA and ionomycin with Brefeldin A and were analyzed by flow cytometry (n=3). **d** PBMCs were isolated from healthy human volunteers and stimulated with PMA and ionomycin in the presence or absence of cyclocreatine (n=2-5). RNA was isolated and evaluated by qPCR relative to actin. Statistical differences were analyzed by 2way ANOVA (**a,b,c**) and unpaired T test (**d**)P&lt;0.01, *** P&lt;0.001

Since T cells are among the primary tissue IFN-γ producers, we evaluated relative number of CD3^+^ cells in DSS treated colon tissue and found equal overall numbers of CD3^+^ cells both visually and when quantified per high power field (Supp Figure 1B and 1C, p >0.05). This data suggested that differences in T cell numbers did not contribute to the differences in cytokine expression.

### Loss of CK results in reduced IFN-γ production in response to stimulation

To evaluate the intrinsic loss of IFN-γ in CKdKO cells independent of DSS stimulation, we isolated lymphocytes from control and CKdKO mice. We found that in response to PMA/Ionomycin stimulation, CKdKO peripheral blood lymphocytes produced significantly less IFN-γ by ELISA (Fig 2B, p<0.01) compared to control cells. In order to identify the contribution of specific cell types in the loss of IFN-γ, splenocytes were isolated and stimulated with PMA/Ionomycin to determine the ability of the cells to produce IFN-γ by flow cytometry. As shown in Figure 2C, this analysis revealed an IFN-γ production defect in both CD4^+^ (p<0.001) and CD8^+^ T cells (p<0.01) with no evidence of differences in peripheral T cell numbers.

To determine if the IFN-γ production defect translates to human cells, we made use of the creatine mimic cyclocreatine with human peripheral blood mononuclear cells (PBMCs).

Cyclocreatine partially inhibits both CRT and CK activity by competing with creatine for transport and phosphorylation^21^. We treated human PBMCs with stimulation (PMA/ionomycin) in the presence of cyclocreatine and identified an almost complete loss in IFN-γ protein secretion (Fig 2D, p<0.001).

### Addback of IFN-γ partially rescues CKdKO DSS susceptibility

To determine the relative contribution of IFN-γ to the observed increase in DSS colitis disease severity within CKdKO mice, we administered exogenous IFN-γ by intraperitoneal (i.p.) injection during the cycle of DSS colitis. We found a significant improvement in weight loss scores associated with i.p. IFN-γ administration as compared to PBS controls (Fig 3A, p<0.01). No differences in overall disease activity or colon length were observed with exogenous IFN-γ (Fig 3B and 3C, p >0.05). Further analysis revealed a notable improvement in histologic scores with largely preserved crypt architecture and decreased inflammatory infiltrate in colonic tissue of CKdKO mice following administration of IFN-γ (Fig 3D and E, p<0.05). The disparity in effect between disease activity index and histology may be due to improved healing which did not yet affect disease symptoms at the given timepoints. Although exogenous IFN-γ did not completely normalize disease activity in CKdKO mice, there was significant protection to indicate that the CK driven loss of IFN-γ contributes in part to the disease susceptibility.

**Figure 3.**
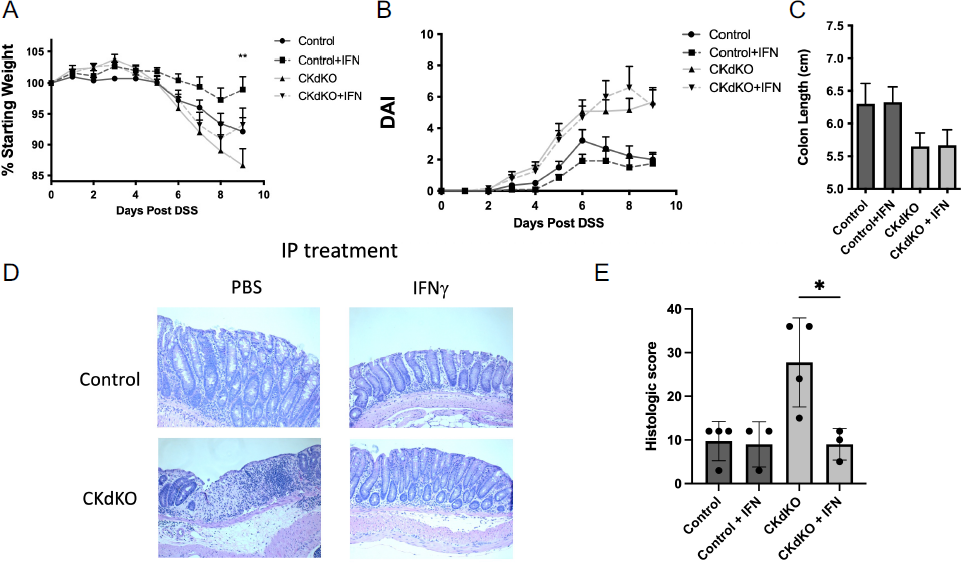
CKdKO and control mice were exposed to 1.5% DSS in their drinking water for 5 days with the addition of IP injections of IFN-γ or PBS followed by a 4-day recovery period. **a** Weights and **b** Disease Activity Index (weight change, stool consistency and bleeding score) were obtained daily. **c** Colon length was measured from cecum to distal colon. **d, e** H & E of colonic tissue were compared and scored (see methods section). n=12-14 mice per group for **a, b, c** and n=4 for **e**. Statistical differences were analyzed by 2way ANOVA (**a, b)**, unpaired t test (**c**) and one-way ANOVA (**e**). *P&lt;0.05, ** P&lt;0.01

### Hypoxia inducible factor (HIF) is stabilized in CKdKO splenocytes

To identify the mechanism(s) by which CKdKO lymphocytes produce less IFN-γ, we utilized an unbiased transcription factor array. We conducted this evaluation comparing unstimulated splenocytes isolated from control and CKdKO mice. Interestingly, CKdKO splenocytes showed a >40-fold increase in hypoxia inducible factor (HIF) compared to controls (Supp Fig 2). HIF is widely accepted as a central regulator of tissue metabolism^22^ and has been shown to be important in the resolution of acute inflammation in the mucosa^14^. It is also notable that we have previously identified HIF as a regulator of CK expression in intestinal epithelial cells^11^. HIF has not previously been shown to regulate CK expression in lymphocytes, outside the setting of malignancy.

To confirm the stabilization of HIF in CKdKO lymphocytes, we isolated splenocytes from control and CKdKO mice and evaluated for HIF-1α protein stabilization in the presence and absence of PMA/Ionomycin stimulation. Consistent with resulted from the protein array, HIF-1α was profoundly increased in CKdKO splenocytes at baseline and in the presence of stimulation (Fig 4A).

**Figure 4.**
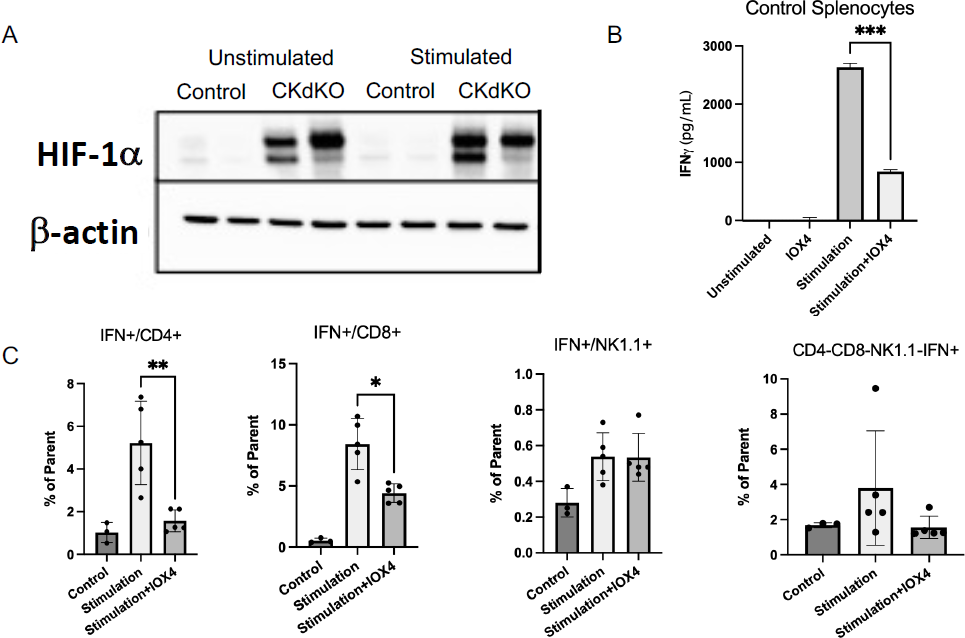
**a** Splenocytes were isolated from wildtype and CKdKO mice and incubated in the presence or absence of PMA and ionomycin. Protein was then collected from the cells and HIF-1α was assessed by western blot relative to actin. **b** Isolated wildtype splenocytes were treated with HIF stabilized IOX4 in the presence or absence of PMA, ionomycin and α-CD3 activation. Supernatants were obtained evaluated by ELISA. **c** Splenocytes stimulated with PMA, ionomycin or α-CD3 in the presence or absence of IOX4 were evaluated by flow cytometry for intracellular IFN-γ in CD4^+^, CD8^+^ and NK1.1^+^ populations. n=3-5. Statistical differences were analyzed by unpaired t test (**b**) and one-way ANOVA (**c**). * P&lt;0.05, ** P&lt;0.01, *** P&lt;0.001.

### HIF stabilization reduces IFN-γ production by CD4^+^ and CD8^+^ T cells

Based on the observations that deletion of CK resulted in decreased IFN-γ production by accompanied by HIF-1α stabilization, we next sought to determine if, independent of CK expression, HIF stabilization results in reduced IFN-γ production. To do this, we utilized a pharmacological approach incorporating the HIF stabilizing agent (prolyl hydroxyly inhibitor) IOX4^23^. As shown in Figure 4B, IOX4 significantly reduced IFN-γ production by anti-CD3-stimulated splenocytes (p<0.001). We confirmed these findings by intracellular IFN-γ flow cytometry staining. CD4^+^ (p<0.01) and CD8^+^ (p<0.05) splenocytes produced less intracellular IFN-γ with IOX4 treatment (Fig 4C). Notably, no change in intracellular IFN-γ was observed in NK1.1^+^ NK cells or the remaining splenocytes (p>0.05). These data suggest that HIF-1α stabilization is sufficient to induce the loss of IFN-γ observed in CKdKO mice during DSS colitis.

## Discussion

The mucosal tissue environment of the intestine is complex, involving interactions between the immune system, the epithelium and the constellation of microorganisms (microbiome) present in the lumen. This tissue environment changes during episodes of inflammation to include profound shifts in metabolism. In work dating back to the 1980’s, Roediger et. al. proposed the “starved gut” hypothesis wherein they identified that IBD patient colonocytes are deficient in short chain fatty oxidation to the extent that energy deficiency becomes a dominant phenotype in these patients^24–27^. In the ensuing 40 years, additional findings have supported this finding, including inflammation-associated shifts in the microbiome and multiple metabolites, that perturb tissue energetics and cellular metabolism ^28^.

Previous studies made the important observation that creatine metabolism and creatine transport are abnormal in IBD^11, 17^. To build on these studies, we sought to define the influence of global deletion of CKB and CKMit on colitis outcomes in a murine model. We found that CKdKO mice develop significantly more severe colitis in the DSS colitis model. This finding is important given known CK dysfunction in the mucosa of IBD patients and may suggest that the loss of CK expression could contribute to continued disease activity.

Interestingly we found that despite worsened disease activity, there was loss of several cytokines traditionally considered to be proinflammatory. Most significant was the loss of IFN-γ in the colonic tissue and histologic improvement with IFN-γ add back in DSS colitis. This is intriguing, as in addition to the proinflammatory role of IFN-γ, there is an important role for IFN-γ in disease resolution by blocking protease activities and blocking immune cell infiltrate^19,20^. While DSS studies in IFN-γ knockout mice results in protection and argues for an inflammatory role for IFN-γ in the digestive tract^29^, these prior studies and our findings suggest that IFN-γ titration is vital to achieving inflammation as well as inflammatory resolution. Interestingly in a number of different immune cell populations including mucosal-associated invariant T cells^30^ and mononuclear cells^31^ isolated from IBD patient samples produce less IFN-γ than healthy controls again arguing for dysregulation in IFN-γ in IBD disease pathogenesis.

In addition to the loss of IFN-γ seen in CKdKO DSS tissue, we also identified reductions of other key proinflammatory cytokines such as IL-22, IL-6 and IL-17F. IL-22 is protective in the mucosal environment by inducing proliferative pathways in the epithelium and thus makes sense for causing increased colitis susceptibility. Although we found both a significant HIF-1α stabilization in the peripheral lymphocyte and loss of IL-22 in the mucosal tissue, previous research has identified a positive relationship between HIF stabilization and IL-22 production by CD4+ T cells^32^. These findings suggest that other pathways are also likely involved in the regulation of IL-22 in the setting of CK loss and warrants further study. Interestingly, HIF stabilization by DMOG treatment of DSS treated mice resulted in disease protection and loss of

IL-6 production^33^ suggesting that HIF may also contribute to regulation of IL-6 with the loss of CK activity. Similarly, to IFN-γ, IL-6 has both pro and anti-inflammatory attributes as IL-6 producing lymphocytes were found to be vital to epithelial barrier and health^34^. A recent publication identified a close relationship between IL-17A and to a lesser degree IL-17F is vital to epithelial wound healing by HIF-1α activation^35^. This could contribute to the worsened disease status in CK deficient mice with reduced IL-17F production. IL-17 expression has been linked to HIF-1α activity, with HIF knockdown mice producing less IL-17A and developing reduced colitic disease in a RAG lymphocyte transfer model^36^. This would suggest that HIF-1α stabilization should increase IL-17 production, which is the opposite of what we have demonstrated, arguing for an alternate mechanism for this cytokine loss.

At present, we do not know the mechanism(s) of HIF stabilization in CKdKO lymphocytes. It is widely known that HIF functions as a global metabolic sensor ^37^ and the shifts in intracellular metabolism with loss of CK likely reflects a stress response to the extent that HIF is stabilized. Nonetheless, the association of basal HIF stabilization in CKdKO with IFN-γ implicates HIF in the repression of the IFN-γ gene. It is notable that HIF functions as both a transcriptional activator and a transcriptional repressor, where gene expression profiles have shown a nearly equal number of induced and repressed genes in vascular endothelial cells^38^.

Interestingly, our own examination of the murine and human IFN-γ promoters identified classic HIF response elements (sequence 5’-RACGTG-3’)^39^, providing the distinct possibility that HIF is a direct transcriptional repressor of IFN-γ.

Although IFN-γ add-back in part reversed the severe phenotype in CKdKO mice, it did not completely resolve the increases susceptibility caused by CK loss. This is likely due to other impacts of CK loss in the mucosal environment including changes to the cytokine profile noted above. Given the central role of the creatine pathway in energy regulation, we believe that CK is likely impacting other cellular activities in the intestine. Notably our prior research identified intestinal epithelial dysfunction in the setting of CRT loss^17^. It is reasonable to think that CK loss could similarly impair epithelial cell wound healing and barrier function although these studies remain to be completed.

In summary, CK activity in the intestinal mucosal important is protective in the setting of DSS colitis. This protective effect is in part due to regulation of HIF-1α stabilization which we found to have a suppressive effect on IFN-γ production. Absence of IFN-γ resulted in increased tissue injury as evidenced by the protective influence of IFN-γ add back experiments. In total, these studies confirm that the creatine pathway is a fundamentally important immunologic regulator in the intestinal mucosal environment and loss of creatine regulation may contribute to disease.

## Methods and Materials

### Mouse lines

All wild type littermates that were all bred in house. Animals were maintained and handled according to protocols approved by the Institutional Animal Care and Use Committee (IACUC). Original breeding pairs of CKdKO mice were obtained Dr. Be Wieringa, Radboud University, Nijmegen, Netherlands^40^ and bred in house for the studies described here.

Undisturbed, these mice harbor no outward phenotype other than some learning deficits and acoustic startle reflexes.

### DSS colitis

Male and female mice were subjected to DSS at 8-10 weeks of age. From day 0, mice received drinking water with 1.5% DSS (molecular weight 36,000–50,000; MP Biomedicals) to induce acute colitis, or H_2_O as a vehicle control. After 5 days of treatment, mice were allowed to recover for 2 days on regular drinking water before sacrifice. A disease activity index (DAI) score was assessed daily to evaluate the development of colitis based on the parameters of weight loss compared to initial weight, stool consistency, and rectal bleeding^18^. Evaluation was performed by two researchers who were blinded to the experimental groups. Scores were defined as weight loss: 0 (0%), 1 (1-5%), 2 (5-10%), 3 (11-20%), and 4 (>20%); stool consistency: 0 (well-formed pellets), 2 (pasty, semi-formed pellets), and 4 (liquid stools); and rectal bleeding: 0 (no blood), 2 (hemoccult positive), and 4 (gross bleeding). Colon lengths were measured at time of sacrifice, and tissue collected for immunofluorescence, RNA, and protein analyses.

Histological scoring of paraffin-embedded tissues were as described before ^41^. The three independent parameters measured were severity of inflammation (0-3: none, slight, moderate, severe), extent of injury (0-3: none, mucosal, mucosal and submucosal, transmural), and crypt damage (0-4: none, basal 1/3 damaged, basal 2/3 damaged, only surface epithelium intake, entire crypt and epithelium lost). The score of each parameter was multiplied by a factor reflecting the percentage of tissue involvement (x1: 0-25%, x2: 26-50%, x3: 51-75%, x4: 76-100%) and all numbers were summed. Maximum possible score was 40. For IFN-γ add back experiment, recombinant murine IFN-γ (Biolegend) or PBS was administered by IP injection at 0.25 mg/kg on days 3-8 of DSS exposure. Dosing regimen was based on previous work^42^.

### Cytokine analysis

Proinflammatory cytokines in the terminal third of murine colon samples were harvested, snap frozen in liquid nitrogen and stored at −80°C until further analysis. Tissue was thawed in sample buffer and solubilized using a tissue homogenizer. Cytokines were measured using the murine proinflammatory assay (Mesoscale, Gaithersburg, MD, USA).

### CD3 Immunofluorescence analysis

Colonic tissue was harvested from DSS colitis experiments and fix 10% buffered formalin fixed prior to paraffin embedding. Tissues were deparaffinized and stained with anti-CD3 primary antibody (Abcam, 1:100 dilution) followed by anti-rabbit FITC secondary (Biolegend) and counter stained with DAPI. Resulting images were analyzed for numbers of CD3 positive T cells per high power field using Image J. A minimum of 10 fields per image were included in the analysis.

### Mouse peripheral blood lymphocyte and splenocyte isolation

Briefly, whole blood was collected via cardiac puncture into syringes containing anticoagulant (K_2_EDTA at 1.8mg/ml blood). Blood was gently layered over double-density Histopaque gradients (1119 / 1083) and centrifuged at 700 x *g* in a swinging bucket rotor centrifuge for 30min without brake. The resulting PBL layer was collected, and residual red blood cells lysed and used in conditions described within the text. Mouse splenocytes were obtained by excising the spleen from euthanized mouse. Splenocytes were isolated by extruding tissue through a 70-um filter into RPMI media (RPMI +0.1% BME, 1% Penicillin/streptomycin, 10% FBS). Cells were spun at 1000xg, and supernatant removed. Cells were then treated with RBC lysis buffer (0.15 M NH2Cl, 10 mM KHCO3, 0.1mM EDTA) for 1 minute at room temperature and then washed with 10 ml of RPMI. Cells could then be counted and used in subsequent experiments.

### Human PBMC isolation

Human PBMCs were isolated from whole venous blood of healthy volunteers (IRB# 06–0853). Whole venous blood was collected in syringes containing anticoagulant (K_2_EDTA at 1.8mg/ml blood). Blood was gently layered over Histopaque 1077 and centrifuged at 700 × *g* in a swinging bucket rotor centrifuge for 30min without brake. The resulting PBMC layer was collected, and residual red blood cells lysed as previously described. PBMCs were washed with ice-cold HBSS-(w/out CaCl_2_ or MgCl_2_), counted and used for qPCR. Human PBMCs were collected and stimulated with PMA/ionomycin (10 ng/ml and 1µg/ml, respectively) in the presence of cyclocreatine (5mM final concentration) or vehicle control (media only) for 5 hours. RNA was extracted using Trizol (Sigma) and evaluated for IFN-γ (Primer sequence: forward : TCGGTAACTGACTTGAATGTCCA; reverse : TCGCTTCCCTGTTTTAGCTGC).

### IFN-γ ELISA

ELISA experiments were conducted using mouse peripheral blood lymphocytes and mouse splenocytes. Mouse cells were incubated in the presence of PMA/ionomycin (20 ng/ml, 1ug/ml) and plate bound anti-CD3 (500 ug/ml) for 4 hours. In later experiments, the HIF stabilizer IOX4 (5 uM, MedChem Express) was added during the stimulation exposure.

Supernatants were collected and evaluated using standard IFN-γ ELISA Max Deluxe Mouse (Biolegend).

### Flow cytometry

Splenocytes from CKdKO and control mice were isolated as above, then stimulated with PMA/ionomycin (20 ng/ml, 1ug/ml) and plate bound anti-CD3 (500 ug/ml) in the presence of Brefeldin A (1:1000, Biolegend) for 4 hours. In subsequent experiments IOX4 (5uM) was added for HIF stabilization. Cells were then stained for surface stains: NK1.1, CD8, CD4 (Biolegend) as well as a live/dead marker (Invitrogen) for 30 minutes at 4C in FACS buffer (5% FBS in PBS).

Cells were washed in Flow buffer and then fixed and permeabilized using FoxP3/Transcription Factor Staining Buffer Set (eBioscience). Intracellular staining of IFN-γ (Biolegend) was conducted for 30 minutes at room temperature. Cells were washed and held at 4C until run on BD FACS Canto II machine and analyzed by FlowJo.

### Transcription factor array

Unstimulated splenocyte nuclear samples were arrayed on a TF Activation Profiling Plate-I monitoring 48 different transcription factors (Signosis, Santa Clara, CA). Analysis was performed as recommended by the manufacturer.

### HIF Western blot

Splenocytes were isolated from CKdKO, and control mice as previously described and then treated with PMA/Ionomycin (20 ng/ml, 1ug/ml) or no treatment for 4 hours. Cells were spun down and pellets dissolved in 1x SDS-PAGE Sample Buffer (Bio-Rad) containing 100 mM DTT, 5 mM EDTA, and 1x HALT Protease Inhibitor Cocktail. Samples were sonicated and loaded on 4-20% Mini-PROTEAN TGX Precast Protein Gel (Bio-Rad). Samples were transferred to 0.2um PVDF membranes using a Bio-Rad RTA kit and using a Bio-Rad Trans-Blot Turbo instrument.

Blots were blocked with 5% nonfat dry milk (Bio-Rad) in TBS-T. Blots were incubated with primary antibodies mouse anti-HIF1A (BD 610959, 1/500) and rabbit anti-ACTB (Abcam ab8227, 1/10,000) overnight at 4C. Blots were washed 4 times for 10 minutes with TBS-T then incubated for 1 hour at room temperature using secondary antibodies goat anti-mouse (MP Bio 0855550, 1/10,000) or goat anti-rabbit (MP Bio0855676, 1/10,000). Blots were again washed 4 times for 10 minutes with TBS-T and developed using Clarity Max ECL reagent. Imaging was conducted using Bio-Rad ChemiDoc MP instrument.

### Statistical analysis

All raw and calculated data are expressed as mean±standard error of the mean (SEM) of n observations, n being the number of biological replicates, and analyzed using Prism 5.0 (GraphPad San Diego, CA, USA). Changes in transcript and protein levels were compared using Student’s unpaired t-test or one way ANOVA with Newman-Keuls post-hoc test where appropriate. A p-value <0.05 was considered significant.

## Data Availability

Data sharing not applicable to this article as no relevant datasets were generated or analyzed during the current study.

---

**Supplemental Figure 1.**
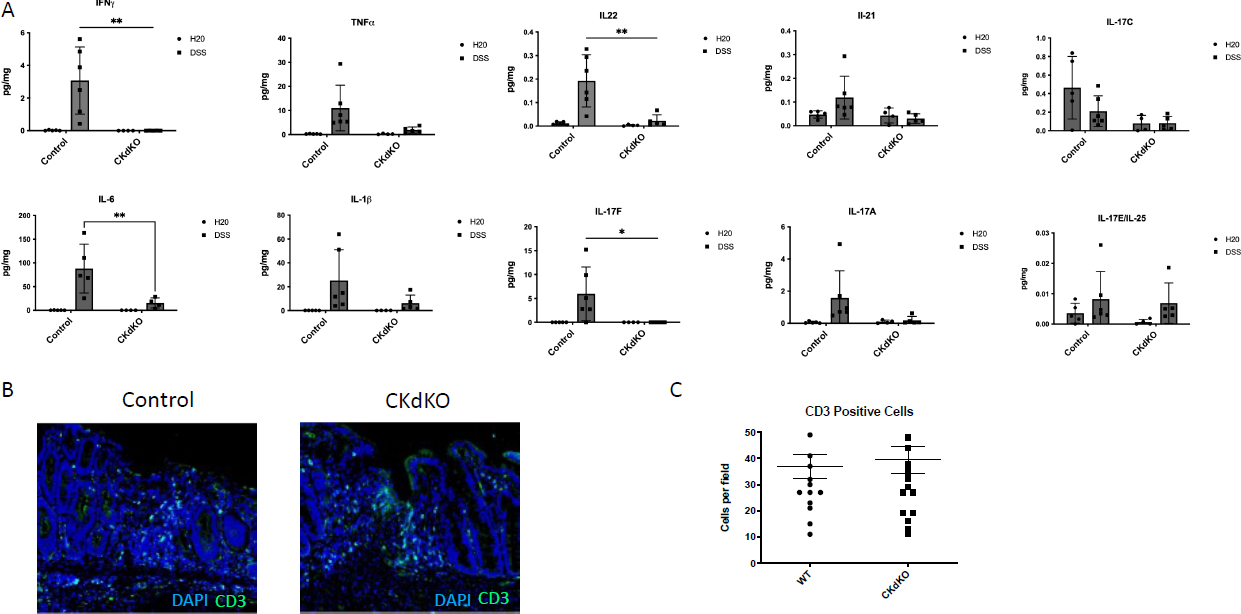
**a** Complete colonic tissue mesoscale protein analysis from DSS colitis experiment in wildtype and CKdKO mice. **B** DSS colitis colonic tissue was analyzed for CD3 cell infiltrate. **c** Quantification of CD3+ cells/high powered field. n=6 (**a**) and n=6 (**c**). Statistical differences were analyzed by 2way ANOVA (**a**) and unpaired t test (**b**). * P<0.05, ** P<0.01

**Supplemental Figure 2.**
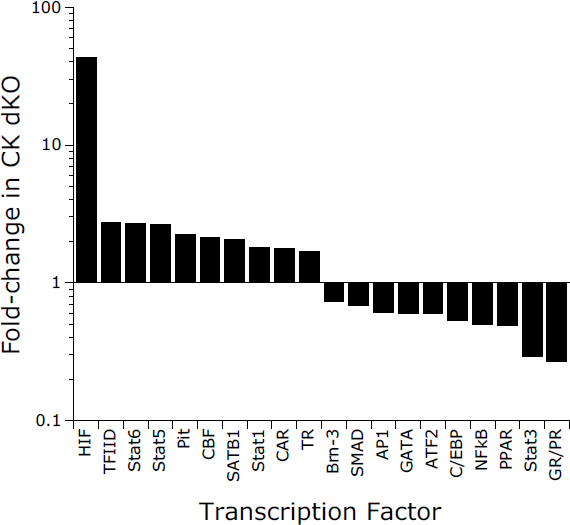
Splenocytes isolated from wildtype and CKdKO mice were evaluated by transcription factor array with the most significantly different transcription factors displayed.

## Acknowledgements

Thank you to the members of the Colgan Lab and the University of Colorado Mucosal Inflammation Program for feedback on this project. This work was supported by grants from the National Institutes of Health, the Veterans Administration and by the Crohn’s and Colitis Foundation.

## Author Contributions

CHTH, JML and SPC conceived the study and initiated the study design, conducted experiments and analysis. CHTH wrote the manuscript with contributions from JML and SPC. ASD and EMM contributed to the study implementation and analysis. Review and editing of final manuscript were conducted by all authors.

## Competing interests

The authors have no financial competing interests to disclose.

## Notes

### Competing Interest Statement

The authors have declared no competing interest.

